# Diffusion MRI free water is a sensitive marker of age-related changes in the cingulum

**DOI:** 10.1101/867606

**Authors:** Manon Edde, Guillaume Theaud, François Rheault, Bixente Dilharreguy, Catherine Helmer, Jean-François Dartigues, Hélène Amieva, Michèle Allard, Maxime Descoteaux, Gwénaëlle Catheline

**Affiliations:** EPHE, PSL, F-33000, Bordeaux, France; CNRS, INCIA, UMR 5287, F-33000 Bordeaux, France; Université de Bordeaux, INCIA, UMR 5287, F-33000 Bordeaux, France; Université de Bordeaux, Inserm, Bordeaux Population Health Research Center, UMR 1219, F-33000 Bordeaux, France; CHU de Bordeaux, Bordeaux, France; Sherbrooke Connectivity Imaging Lab (SCIL), Université de Sherbrooke, Sherbrooke, QC, Canada

**Keywords:** aging, tractography, free water, white matter hyperintensity, cingulum

## Abstract

Diffusion MRI is extensively used to investigate changes in white matter microstructure. However, diffusion measures within white matter tissue can be affected by partial volume effects due to cerebrospinal fluid and white matter hyperintensities, especially in the aging brain. In previous aging studies, the cingulum bundle that plays a central role in the architecture of the brain networks supporting cognitive functions has been associated with cognitive deficits. However, most of these studies did not consider the partial volume effects on diffusion measures. The aim of this study was to evaluate the effect of free water elimination on diffusion measures of the cingulum in a group of 68 healthy elderly individuals. We first determined the effect of free water elimination on conventional DTI measures and then examined the effect of free water elimination on verbal fluency performance over 12 years. The cingulum bundle was reconstructed with a tractography pipeline including a white matter hyperintensities mask to limit the negative impact of hyperintensities on fiber tracking algorithms. We observed that free water elimination improved the sensitivity of conventional DTI measures to detect associations between tissue-specific diffusion measures of the cingulum and changes in verbal fluency in older individuals. Moreover, free water content measured along the cingulum was independently strongly associated with changes in verbal fluency. These observations suggest the importance of using free water elimination when studying brain aging and indicate that free water itself could be a relevant marker for age-related cingulum white matter modifications and cognitive decline.

## 1. Introduction

Aging is associated with widespread brain structural modifications including neurodegeneration of white and grey matter [1–3]. In the last few years, diffusion tensor imaging (DTI) has been widely used to indirectly explore microstructural properties of white matter and constitutes a sensitive technique to describe age-related white matter microstructural changes [4,5]. Parameters quantified by DTI such as Fractional Anisotropy (FA), Mean Diffusivity (MD) and Radial Diffusivity (RD) can be used to indirectly infer changes in axonal integrity and myelination [6]. In older adults, previous DTI studies reported a decrease in FA and an increase in MD and RD in major white matter tracts [6–8] that correlated with cognitive impairment [9–12].

However, diffusion measures within brain tissues can be affected by partial volume effects due to cerebrospinal fluid [13], especially in aging individuals with atrophied brains and ventriculomegaly [14–17]. When voxels contain cerebrospinal fluid (e.g. partial volume), diffusion measures can be biased towards a pattern of high diffusivity (MD, Axial Diffusivity (AD), RD) and reduced FA [18,19]. This effect has been particularly observed in the fornix and the corpus callosum because of their close proximity to periventricular regions [18,20,21]. To overcome this problem, the Free Water Elimination (FWE) method was used to differentiate the diffusion properties of brain tissues such as white matter bundles from the surrounding free water composed of cerebrospinal fluid [13,22]. Elimination of extracellular free water contamination improves the sensitivity and specificity of most measures derived from conventional DTI [16,22–24]. Although such correction has been used in pathological conditions [25–30], the majority of diffusion MRI studies do not consider free water effects of aging. Aging is associated with grey and white matter tissue loss, and the resulting enlargement of interstitial space could lead to increase in free water [31–34]. In addition, free water content in white matter fibers was recently associated with the presence of white matter hyperintensities (WMHs) in clinical studies [28,35–38]. In older adults, the prevalence and severity of WMH burden increases after age 60 [39,40]. Previous studies reported that the presence of WMH lesions was not only associated with age-related white matter changes evaluated with DTI (Maillard et al., 2014; Maniega et al., 2015; Pelletier et al., 2015) or tractography (Reginold et al., 2016, 2018; Svärd et al., 2017), but also with cognitive impairments mainly in memory and executive functions [5,46,47].

The cingulum bundle is one of the most studied white matter tracts running from the anterior to the posterior part of the brain that interconnects frontal, parietal, and temporal areas (Catheline et al., 2010; Jones et al.,2013; Bubb et al., 2018). Due to this central location, the cingulum plays a central role in white matter architecture and functional networks that support and contribute to optimal cognitive function (Bubb et al., 2018). In previous studies, changes within this bundle due to aging were correlated with executive and memory deficits (Catheline et al., 2010; Ezzati et al., 2016; Kantarci et al., 2011; Metzler-Baddeley et al., 2012; Seiler et al., 2018; Sexton et al., 2010). However, these observations still remain controversial considering that most of these studies did not consider CSF partial effect due to atrophy or WMH burden. Recent tractography studies consistently observed that atrophy-related partial volume effects [16,18] and age-related ventricular enlargement [15] or WMH burden (Reginold et al., 2016) affect diffusion measures of the cingulum in healthy older individuals. In this regard, Reginold and colleagues reported higher RD in cingulum tracts that crossed WMH than those outside WMH regions (Reginold et al., 2016; see also Kurki & Heiskanen, 2018). Atrophy and WMH burden might therefore impact cingulum diffusion measures suggesting a source of significant bias in studies exploring the integrity of this bundle and its association with cognitive functions in aging.

In this study, we hypothesized that free water elimination would provide more accurate white matter diffusion measures of the cingulum, which in turn would be better correlated with verbal fluency changes in 68 elderly individuals. To do so, the cingulum bundle was reconstructed with a tractography pipeline that includes a WMH mask to limit the negative impact of hyperintensities on fiber tracking algorithms. We first described the effect of FW-correction on diffusion measures of the cingulum derived from the conventional tensor DTI model. Then, we explored the association between FW-corrected diffusion measures and cognitive changes as evaluated using the Isaacs Set Test over 12 years. The Isaacs Set Test was chosen since it specifically assesses semantic memory and executive functioning.

## 2. Methods

### 2.1. Dataset

Participants were selected from the Bordeaux sample of the Three-City study [55], an ongoing longitudinal multicenter population-based elderly cohort designed to evaluate risk factors for dementia. The study protocol was approved by the ethics committee of Kremlin-Bicêtre University Hospital (Paris, France), and all participants provided written informed consent. At baseline, subjects were non-institutionalized, randomly recruited from electoral lists, and were older than 65 years. Since the 1999-2000 baseline inclusion, an extended neuropsychological assessment was administered by trained psychologists at each follow-up occurring at 2, 4, 7, 10 and 12 years. An MRI scan was performed at the 10-year follow-up for every subject. All 239 subjects were screened, and individuals were excluded if they had dementia (n=8), brain pathologies (n=27) or if MRI images were either unavailable (n= 42), distorted by artefacts (n=24), or failed pre-processing (n=15). In addition, cingulum reconstruction failed in 16 subjects and tractography quality control revealed suboptimal data in 19 additional subjects. Out of the 239 initially screened subjects, 68 subjects were included in the study. All participants were right-handed and had a Mini Mental State Examination (MMSE) score greater than or equal to 24.

### 2.2. Cognitive and clinical variables

The Isaacs Set Test (IST, Isaacs & Kennie, 1973) chosen for the study consists of a test on verbal fluency, where subjects are asked to cite the highest number of words in four semantics categories. Three scores of the IST were used, corresponding to the number of words cited by the subject at 15-seconds (IST 15s), 30-seconds (IST 30s) and 60-seconds (IST 60s). Verbal fluency was evaluated over 12 years, using a subject-specific slope for each IST score computed using a linear mixed model with time as a continuous variable, random intercept and slope. Over the 12-year follow-up period, the negative mean annual slope indicated a decrease in the number of given words. Some clinical variables were also collected: depressive symptoms evaluated using the Center for Epidemiologic Studies-Depression scale (CESD, Radloff, 1977), vascular risk factors including hypertension as defined in patients with a blood pressure > 140/90 mm Hg or taking anti-hypertensive medication, diabetes as defined in patients with a blood glycemic levels > 7 mmol/l or taking hypoglycemic medication, and body mass index (BMI).

### 2.3. MRI acquisition

MRI scanning was performed using a 3T Achieva system (Philips Medical Systems, The Netherlands) equipped with a 8-channel SENSE head coil. Head motions were minimized by using tightly padded clamps attached to the head coil. Anatomical MRI volumes were acquired using a 3D magnetization-prepared rapid gradient-echo (MPRAGE) T1-weighted sequence with the following parameters: repetition time (TR)/ echo time (TE) = 8.2 ms/3.5 ms, flip angle 7°, field of view (FOV) 256×256 mm^2^, 180 slices of 1 mm of thickness, voxel size 1×1×1 mm^3^. Fluid-attenuated inversion recovery (FLAIR) images were obtained with the following parameters: TR=11000 ms; TE=140 ms, inversion time = 2800 ms, 90-degree flip angle, FOV 230×172 mm^2^, 24 slices of 5 mm of thickness, voxel size 0.72×1.20×5 mm^3^. Diffusion weighted images were acquired using a spin echo single shot EPI sequence composed of one b0 map (b-value=0 s/mm^2^) followed by 21 diffusion gradients maps (b-value=1000s/mm^2^) homogenously spread over a half sphere and 21 opposite directions spread over the opposite half sphere. To increase signal-to-noise ratio, a second series of two b0 and 42 DWI volumes was acquired. Sixty axial slices were acquired with the following parameters: TR= 9700 ms, TE= 60 ms, flip angle 90°, FOV 224×224 mm^2^, 60 slices, no gap and voxel size 2×2×2 mm^3^. All acquisitions were aligned on the anterior commissure-posterior commissure plan.

### 2.4. MRI preprocessing

#### 2.4.1. Diffusion MRI preprocessing

Diffusion MRI (dMRI) images were pre-processed using FMRIB’s Diffusion toolbox [58,59] in order to produce individual FA, MD, AD and RD maps in native space. For each subject, dMRI images were co-registered to the b0 reference image with an affine transformation and were corrected for motion and eddy current distortions. Brain Extraction Tool (BET, Smith, 2002) was applied to eliminate non-brain voxels and resulting dMRI images were denoised using the non-local mean denoising method [61]. To increase signal-to-noise ratio, the two reversed phases were then combined into a dMRI image using FSL tools. Finally, the fiber Orientation Distribution Functions (fODF) map was computed using the spherical constrained deconvolution (Descoteaux et al., 2007; Tournier et al., 2007; Theaud et al., 2019). Visual quality check was performed and did not reveal any processing failure for the included subjects.

#### 2.4.2. White Matter Hyperintensity segmentation

White matter hyperintensity volumes were automatically segmented by the lesion growth algorithm implemented in the Lesion Segmentation Tool (LST, v2.0; Schmidt et al., 2012) of SPM12. For each participant, the T1-weighted image was used to generate partial volume estimation and three tissue probability maps (grey matter, white matter and cerebrospinal fluid). Then, each FLAIR image was co-registered on the corresponding T1 image to compute lesion belief maps for the three tissue classes. These maps were finally summed up and a lesion growth model with a pre-chosen initial threshold (κ = 0.3) was applied to create lesion maps in native space. Visual inspection of the lesion probability maps was performed and manually corrected when inaccuracies were found. Finally, WMH volumes were extracted, normalized by white matter volume and log transformed (total WMH). The volume of WMH within the cingulum was estimated by crossing the WMH mask with the cingulum bundle (cingulum WMH).

#### 2.4.3. Free Water Elimination

Free Water Elimination (FWE, Pasternak et al., 2009) was used to estimate and remove the free water components from diffusion images. To isolate the fractional volume of freely diffusing extracellular water molecules from tissue compartments (i.e. reflecting intracellular processes), FWE fits a bi-tensor model within each voxel: a first one models free water diffusion, defining as an isotropic diffusion, and a second one models the tissue compartment [24]. The isotropic compartment models extracellular water which is characterized by freely and not hindered diffusion. A fixed diffusivity of 3.10^−3^ mm^2^/s (i.e. diffusion coefficient of water at body temperature) is used for this compartment. The fraction of free water content per voxel provided a native free water (FW) map for each subject and varied between 0 to 1. In contrast, the tissue compartment models water molecules close to cellular membranes of brain tissue, which are defined by restricted or hindered diffusion using a diffusion tensor. Thereby, the volume of freely diffusing extracellular water molecules is removed from tissue compartments. Consequently, the FW-corrected measures were expected to be more sensitive and specific to tissue changes than the measures derived from the single tensor model [23,27,29]. The FW-corrected DTI maps were called FA tissue (FAt), RDt, ADt and MDt (Chad et al., 2018).

#### 2.4.4. Cingulum tractography and measures

For each participant, a local tracking using probabilistic algorithm based on fODF maps was performed. The tractography was performed using twenty tracking seeds per voxels included in the white matter mask. The seeding and tracking masks were modified to include WMH masks (see supplementary material). Next, the left and right cingulum bundles were extracted using the RecoBundles approach (Garyfallidis et al., 2018, Figure 1). RecoBundles is a model-based algorithm performing a registration of a Cingulum model on each subject to extract the subject specific cingulum tract from the whole tractogram. Five different Cingulum models were used in this study (RecoBundles). Finally, after visual inspection of cingulum bundles using dmriqc tool (https://github.com/GuillaumeTh/dmriqcpy), 13 subjects were excluded due to the absence of right or left cingulum bundles and 6 subjects were excluded because of miss-segmentation of hippocampal and anterior parts of the cingulum bundle, mainly because of the age range of our population (older than 85 years of age) at the time of MRI sessions. Therefore, the cingulum bundle was considered as a whole bundle and the different subdivisions of the cingulum were not investigated in this study.

**Figure 1.**
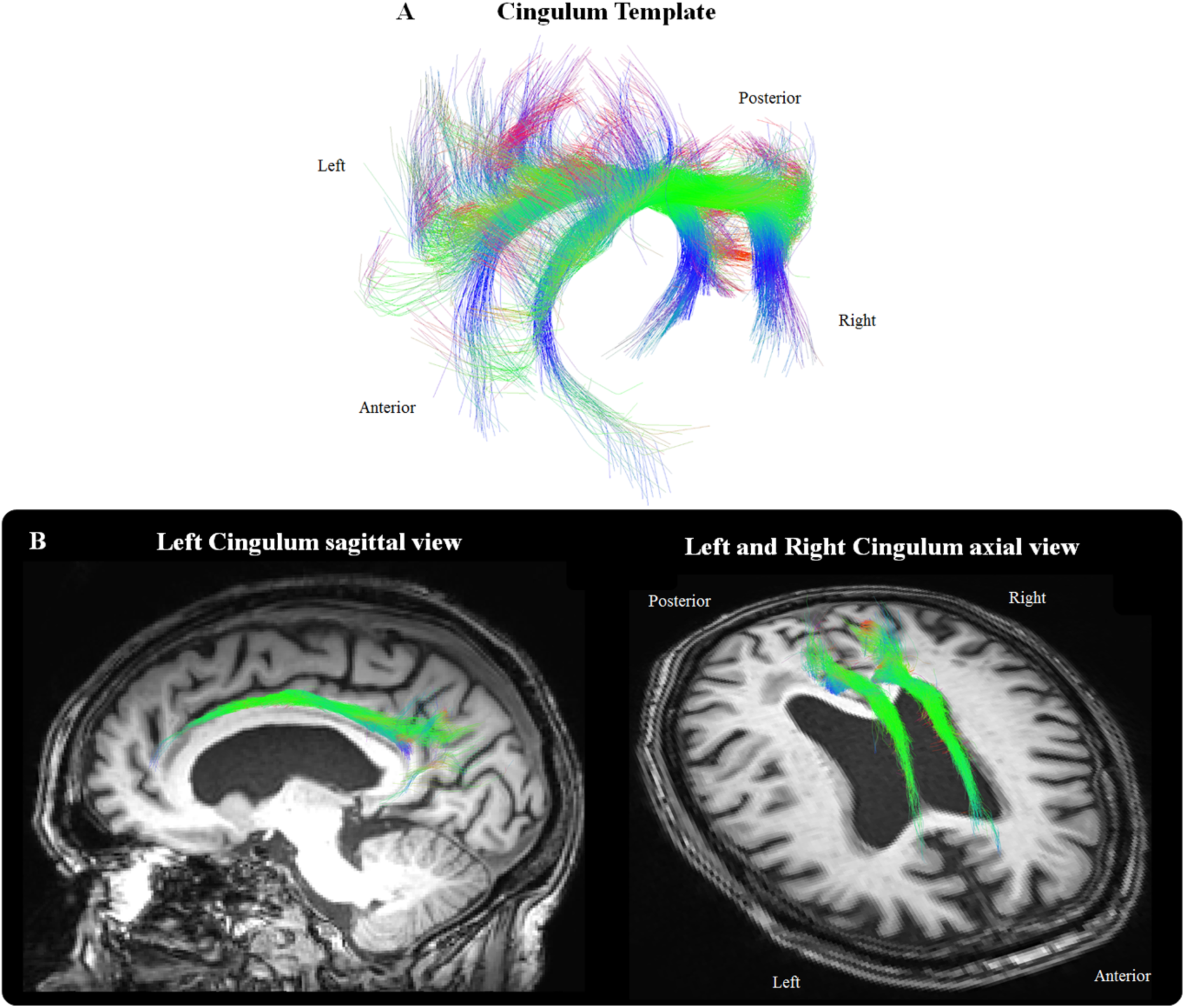
Left and right cingulum templates used to extract the cingulum tract for each participant (A); Example of the cingulum bundle obtained for one subject displayed on the corresponding T1-weighted image (B). Along the fibers, color represents the RGB scale.

Cingulum diffusion measures were extracted before (FA, MD, RD and AD) and after FW-correction (FAt, MDt, RDt and ADt). The diffusion measures without free water correction are defined as conventional DTI measures and reflect the weighted average of all compartments including free water, whereas diffusion measures after correction of free water are defined as FW-corrected measures. Measures of the left and right cingulum were averaged for the analysis.

A percentage of measure changes (% change) for each participant was determined by computing the difference between the measure before and after FW-correction divided by the measure before FW-correction [*i.e*. % change = ((FW-corrected measure – DTI measure)/DTI measure) × 100].

### 2.5. Statistical analysis

Because of non-normality of the continuous and categorical variables, Mann-Whitney U-tests and Spearman’s correlations were used to evaluate the associations between cingulum diffusion measures as well as verbal fluency with continuous variables (age, global cognition score, depressive symptoms score, body mass index and WMH volume) and categorical variables (gender, education level, presence/absence of diabetes, of hypertension) respectively. To compare the effects of FW-correction to conventional DTI on cingulum diffusion measures, a Paired t-Test was performed. Linear regression models were then computed to describe the extent to which the FW-correction affects the association between cingulum diffusion measures and changes in verbal fluency. Diffusion measures were included as independent variables (conventional and FW-corrected FA, MD, AD and RD in separate models) and the verbal fluency slope set as the dependent variable in model adjusted for age and WM volume of the cingulum. Similar additional linear regression models were performed to examine the relationship between free water content and changes in verbal fluency. Total WMH volume was added in previous models to evaluate to the extent to which WMH burden affects the observed associations. In a supplementary analysis, specific WMH burden of the cingulum bundle was added to the models (see S1 Table 3).

A false discovery rate (FDR) multiple-comparison correction method was systematically applied for each analysis. Results were considered significant for p<0.05, FDR corrected [67]. All statistical analyses were performed using IBM SPSS Statistics v.23 software (IBM Corporation, Armonk, NY, USA).

## 3. Results

### 3.1. Sample characteristics

Characteristics of participants are presented in Table 1. Significant age effects on verbal fluency changes were observed for all IST scores (IST 15s, r= -0.212, p=0.022; IST 30s, r= -0.207, p=0.038 and IST 60s, r= -0.252, p=0.019). WMH volumes correlated with age (total WMH: r=0.447, p<0.001; cingulum WMH: r=0.201, p=0.038) and changes in verbal fluency scores (r=-0.218, p=0.038; r=-0.293, p=0.016 and r=-0.328 p= 0.008 respectively). None of clinical variables were related to age, verbal fluency or WMH volumes (p > 0.05). Finally, no association was observed between demographic or clinical variables and cingulum diffusion measures (see S1 Table 1).

**Table 1.** Characteristics of participants,12-year verbal fluency decline and volumetric variables.

| Sample n=68 |  |
| --- | --- |
| *Sociodemographic variables* | Mean ± SD or % |
| Age | 81.2 ± 0.48 |
| Male gender | 36.8% |
| High level of education | 37% |
| *Clinical variables* |  |
| MMSE | 27.3 ± 0.3 |
| CES-D | 8.3 ± 0.9 |
| BMI (kg/m<sup>2</sup>) | 25.2 ± 0.5 |
| Diabetes | 10.3% |
| Hypertension | 35.3 % |
| *Verbal fluency decline* |  |
| IST at 15 seconds | -0.489 ± 0.02 |
| IST at 30 seconds | -0.697 ± 0.04 |
| IST at 60 seconds | -0.924 ± 0.06 |
| *Volumetric variables* |  |
| Total WMH volume ml | 15.9 ± 4.1 |
| Cingulum bundle volume (% TIV) | 0.46 ± 0.06 |
| Cingulum WMH volume (%) | 3.2 ± 0.65 |
| MMSE, Mini Mental State Examination; BMI, Body Mass Index; CES-D, Center for Epidemiologic Studies-Depression scale; IST, Isaacs Set Test; WMH, White Matter Hyperintensity. |  |

### 3.2. Effect of FW-correction on conventional DTI parameters

Cingulum diffusion measures are presented in Table 2. All FW-corrected measures were significantly different from their conventional counterparts (paired t-test, p<0.05 FDR-corrected, Table 2 and Fig 2). After FW-correction, a mean increase of 1.52 % of FAt and a mean decrease of 1.61 % of MDt, 2.5% of RDt and 1.08% of ADt were observed.

**Table 2.** Cingulum diffusion measures before (conventional DTI) and after FW-correction and % of change between both measures for each group

| Cingulum diffusion measures |  |  |  |  |  |
| --- | --- | --- | --- | --- | --- |
| | DTI<br>mean ± SD | FW-corrected<br>mean ± SD | t | p-FDR | % Change<br>mean ± SD |
| FA | 0.543 ± 0.049 | 0.567 ± 0.054 | -3.367 | 0.019* | 1.52 ± 0.50 |
| MD (10<sup>-3</sup> mm<sup>2</sup>/s) | 0.768 ± 0.0041 | 0.755 ± 0.0034 | -3.196 | 0.024* | -1.61 ± 0.22 |
| RD (10<sup>-3</sup> mm<sup>2</sup>/s) | 0.506 ± 0.0047 | 0.493 ± 0.0049 | -5.315 | 0.010* | -2.50 ± 0.19 |
| AD (10<sup>-3</sup> mm<sup>2</sup>/s) | 1.29 ± 0.0683 | 1.27 ± 0.0638 | -2.975 | 0.044* | -1.08 ± 0.34 |
| FW | - | 0.141 ± 0.017 | | | |

\* $p < 0.05$ FDR corrected,
Paired Student's t-test

**Table 3.** Correlations between cingulum diffusion measures and verbal fluency score (IST) before (conventional DTI) and after FW-correction

|  |  | DTI |  |  |  |  |  |
| --- | --- | --- | --- | --- | --- | --- | --- |
|  |  | IST 15s |  | IST 30s |  | IST 60s |  |
| | | $\beta$ | $R^2$ | $\beta$ | $R^2$ | $\beta$ | $R^2$ |
| Model unadjusted for total WMH<sup>a</sup> | FA | 0.015 | 0.001 | 0.047 | 0.002 | 0.067 | 0.004 |
|  | MD | **-0.431\*** | **0.17** | **-0.335\*** | **0.12** | -0.174 | 0.07 |
|  | AD | -0.186 | 0.036 | -0.135 | 0.018 | -0.060 | 0.004 |
|  | RD | -0.114 | 0.013 | -0.128 | 0.016 | -0.129 | 0.017 |
| Model adjusted for total WMH<sup>b</sup> | FA | 0.022 | 0.023 | 0.040 | 0.052 | 0.069 | 0.067 |
|  | MD | -0.267 | 0.020 | -0.218 | 0.010 | -0.147 | 0.09 |
|  | AD | -0.172 | 0.054 | -0.109 | 0.062 | -0.029 | 0.062 |
|  | RD | -0.119 | 0.036 | -0.110 | 0.062 | -0.121 | 0.075 |

|  |  | FW-corrected |  |  |  |  |  |
| --- | --- | --- | --- | --- | --- | --- | --- |
|  |  | IST 15s |  | IST 30s |  | IST 60s |  |
| | | $\beta$ | $R^2$ | $\beta$ | $R^2$ | $\beta$ | $R^2$ |
| Model unadjusted for total WMH<sup>a</sup> | FAt | 0.047 | 0.003 | 0.098 | 0.01 | 0.095 | 0.009 |
|  | MDt | **-0.443\*** | **0.19** | **-0.361\*** | **0.13** | -0.226 | 0.08 |
|  | ADt | -0.198 | 0.04 | -0.134 | 0.018 | -0.061 | 0.004 |
|  | RDt | **-0.293\*** | **0.11** | **-0.248\*** | **0.13** | -0.199 | 0.039 |
|  | FW | **-0.353\*** | **0.12** | **-0.241\*** | **0.10** | -0.196 | 0.061 |
| Model adjusted for total WMH<sup>b</sup> | FAt | 0.035 | 0.044 | 0.072 | 0.061 | 0.071 | 0.07 |
|  | MDt | **-0.401\*** | **0.20** | **-0.341\*** | **0.17** | -0.189 | 0.11 |
|  | ADt | -0.067 | 0.10 | -0.185 | 0.10 | -0.097 | 0.08 |
|  | RDt | -0.224 | 0.11 | -0.213 | 0.11 | -0.173 | 0.09 |
|  | FW | **-0.333\*** | **0.15** | **-0.275\*** | **0.14** | -0.213 | 0.10 |

<sup>a</sup> $\beta$ , standardized coefficient regression adjusted for age and cingulum white matter volume
<sup>b</sup> $\beta$ , standardized coefficient regression adjusted for age, cingulum white matter volume and total WMH volume
$R^2$ , R square value
\* $p < 0.05$ FDR corrected

**Figure 2.**
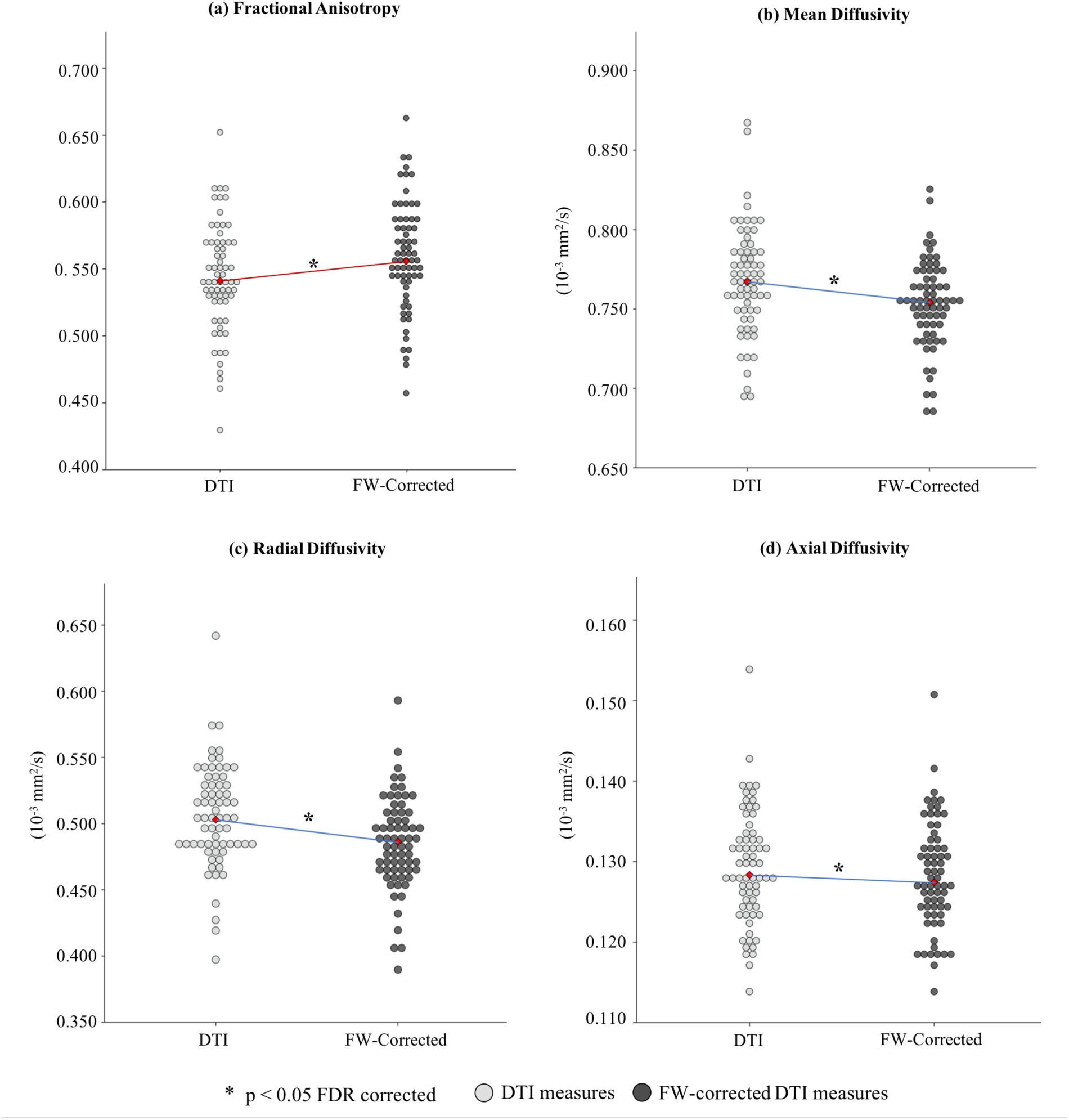
Boxplot charts of the difference in cingulum diffusion measures before (light grey) and after (dark grey) FW-correction for (a) fractional anisotropy, (b) mean diffusivity, (c) radial diffusivity, (d) axial diffusivity. The blue line represents a decrease and the red line represents an increase.

### 3.3. Effect of FW-correction on relationships between cingulum diffusion measures and verbal fluency

#### 3.3.1. Associations with DTI conventional measures

A lower MD value was related to IST decline at 15 and 30 seconds in a model adjusted for age and white matter volume of the cingulum (p < 0.05 FDR-corrected, Table 3). No association was observed with FA, RD and AD.

#### 3.3.2. Associations with FW-corrected measures

After FW-correction, high values of MDt and RDt were strongly associated with IST decline at 15 and 30 seconds in a model adjusted for age and white matter volume of the cingulum (p < 0.05, Table 3). No association was observed with FAt and ADt.

#### 3.3.3. Association with cingulum free water content

High free water content was associated with changes in IST score at 15 and 30 seconds in a model adjusted for age and white matter volume of the cingulum (p < 0.05 FDR-corrected, Table 3).

### 3.4. Impact of WMH burden on the association between cingulum diffusion measures and verbal fluency

In regression models adjusted for total WMH burden, correlations between conventional DTI measures (MD) and IST decline were not significant anymore. In contrast, MDt and free water content correlations were preserved when total WMH volume (Table 3) as well as WMH volume within the cingulum (see S1 Table 3) were added as covariate in regression models (p < 0.05 FDR-corrected, Table 3). In a global model, both diffusion measures remained significantly associated with IST at 15 (FW: β = -0.262 and MDt: β = -0.310) and 30 seconds (FW: β = -0.216 and MDt: β = -0.249, p < 0.05) when adjusted for total WMH volume, indicating that both free water content and MDt independently contributed to the cognitive variances.

## 4. Discussion

In this study, we examined, over a 12-year period, the effect of free water elimination on conventional DTI measures of white matter within the cingulum tract and the effect of such correction on verbal fluency changes in elderly subjects. In a group of 68 older participants, FW-correction significantly impacted all conventional DTI measures and measure following such correction correlated with decline in verbal fluency performances at 15 and 30 seconds. In addition, free water content was also associated with changes in verbal fluency. Finally, in models adjusted for WMH volumes, correlations between MDt and free water content were preserved.

### 4.1. Free water elimination effect

In accordance with recent findings in young adults [22], older adults (Chad et al., 2018) and clinical studies [28,68], FW-correction was associated with an increase in FA, and a decrease in diffusivity measures (MD, RD and AD). These changes in DTI measures after elimination of the free water compartment suggested not only a non-null volume of the extracellular water [6,13,28,69], but also indicate that white matter microstructural changes were less pronounced than previously suggested by conventional DTI measures (Bennett & Madden, 2014; Bennett et al., 2017; Lockhart & DeCarli, 2014; Pelletier et al., 2015).

Based on conventional DTI, our study revealed that only MD showed significant correlations with changes in verbal fluency, especially at 15 and 30 seconds. The elimination of the free water content confirmed such strong association for MDt and revealed additional correlations for RDt that could not be fully observed without considering the free water content. The observed association is in line with previous studies reporting the role of the cingulum bundle in executive functioning in older adults [50,53,73–76]. Even if the underlying neurobiological properties of these parameters remain controversial, a high RDt, without any changes in ADt, has been described as predominantly reflecting myelin changes in animal studies (Song et al., 2003; Sun et al., 2005) and demyelination severity in human post mortem studies on multiple sclerosis (Klawiter et al., 2011; Schmierer et al., 2008). This suggests that changes in verbal fluency in our population might be related to myelin changes within the cingulum, rather than axonal damage (Madden et al., 2012; M. W. Vernooij et al., 2008; Burzynska et al., 2010). Therefore, our results support previous studies reporting that the elimination of free water improves the estimation of tissue-specific indices which were more strongly predictive of cognitive changes than conventional DTI-derived parameters [6,23,26,38,69,77].

### 4.2. Free water content

In our group of elderly participants, a non-null value of free water content was observed, suggesting that despite its distance from the ventricles, some free water is likely present in the extracellular space of the cingulum. This is consistent with recent studies on whole brain white matter reporting a fraction of free water in the elderly (Albi et al., 2017; Chad et al., 2018; Papadaki et al., 2019). In addition, no association between the free water content within the cingulum and CSF volumes was observed in our participants (data not shown). However, we observed that a high content of free water was related to a low cingulum volume, suggesting that in our population the high free water content may be due to atrophy-related processes rather than CSF contamination (Oestreich et al., 2016; Wang et al., 2011). In line with previous older adults and clinical studies, we reported that a high content of free water within the cingulum was strongly associated with changes in verbal fluency performances at 15 and 30 seconds (Bergamino et al., 2017; Ji et al., 2017, 2019; Maillard et al., 2017, 2019). In this regard, in a recent longitudinal study on 224 elderly subjects, Maillard and colleagues showed that an increase in free water was not only associated with reduced performances in executive function assessments but was the strongest predictor of cognitive decline. Taken together, these results support the idea that beyond the elimination of CSF contamination, free water could provide additional structural information that could constitute a physiological index reflecting tissue changes in white matter [27,69,79]. However, the underlying microcellular events accounting for the observed free water content are far from being fully understood; its increase could be related to different pathophysiological processes such as a microvascular degeneration [28,37,80], a reduction in myelin content (Oestreich et al., 2016; Papadaki et al., 2019) or a modulation in the permeability of the blood-brain barrier (Maillard et al., 2017).

### 4.3. White matter hyperintensities burden effect

The present study showed that when adjusting for WMH volumes, the associations between MDt and free water content and verbal fluency performances at 15 and 30 seconds remained significant, suggesting that both could contribute independently to cognitive impairment in aging. In accordance with these results, recent tractography studies reported that cingulum tracts crossing WMH exhibited significant changes in diffusion measures (Reginold et al., 2016), and suggested that these modifications could extend beyond the WMH lesions (Maillard et al., 2014; Promjunyakul et al., 2016; Reginold et al., 2018). Previous DTI studies also reported that WMH were associated with higher diffusion measures in the normal-appearing white matter [41,43,46,81–83]. Taken together, these results suggest that small focal WMH may lead to both local and distant effects that are large enough to impact white matter diffusion properties of the cingulum tract.

Some methodological limitations should be taken into account when interpreting our findings. First, this study was based on a moderate sample size of healthy elderly participants. However, our population exhibited sufficient inter-individual variability in structural measures and cognitive performances to detect associations between these parameters. Second, the cingulum was analyzed as a single entity despite the fact that it consists of a complex structure containing not only long association fibers, but also short tracts connecting adjacent cortices [84–86]. Finally, the current study was based on diffusion MRI data that was acquired with a single b-value. The algorithm used to fit the free water imaging model involved spatial regularization of data which decreases intra-group variability (Pasternak et al., 2009), and may hide subtle spatial features. Even though comparable results using single- and multi-shell acquisitions have been previously reported [87], advanced acquisitions that include multiple b-values could further increase the accuracy of the free water model [69,87].

To conclude, we reported that FW-correction improves the sensitivity to detect associations between tissue specific diffusion measures of the cingulum and changes in verbal fluency in elderly individuals. Moreover, free water content *per se* appears to be a relevant parameter to describe age-related modifications of the white matter and its association with executive functioning. These observations suggest the importance of considering the free water compartment for DTI measures in aging.

## Compliance with ethical standards

### Studies involving human participants

All procedures were conducted with approval from the ethics committee of Kremlin Bicêtre Hospital (Paris, France)

### Informed consent

All participants provided written informed consent for participation in this study.

## Data availability

The data that support the findings of this study are available from the corresponding author upon reasonable request.

## Author contributions

Conceptualization: ME, GC; Formal analysis: ME; Funding acquisition: BD, CH, JFD, HA, MA, GC; Methodology: ME, GT, BD; Software: GT, FR, MD; Supervision: MD, GC; Visualization: ME, GT; Writing - original draft: ME, GT; Writing - review & editing: all authors review the article. All authors provided critical feedback and approved the final manuscript.

