## Supplementary for "Diffusion MRI free water is a sensitive marker of age-related changes in the cingulum"

### Supplementary Material

#### 1. Correction of tractography with white matter hyperintensities (WMH)

In elderly individuals, the presence of WMH can lead to a misclassification during the segmentation process considering that a voxel within WMH could be attributed to grey matter. This misclassification is an issue in tractography since tracking and seeding masks are generated from gray and white matter maps. Therefore, this misclassification can lead to a premature ending of tracks in WMH instead of grey matter. Another important issue of WMH lesions in tractography is the decrease of FA inside WM lesions that could also lead to premature ending of tracks. However, we observed that the principal directions under the lesion mask seem to be preserved and remain coherent and that tractography should be allowed to explore these regions (Figure 1). To determine the effect of WMH on tractography, we produced two whole brain tractograms: a first one without WMH correction and a second one with WMH correction.

The seeding and tracking masks were modified to include the WMH mask, as previously described (Theaud et al., 2017). Briefly, as described in the Methods section, WMH maps were segmented using the Lesion Segmentation Tool (LST, v2.0; Schmidt et al., 2012), visually inspected and registered to the corresponding dMRI images using ANTs registration tool. Finally, WMH maps were added in the inclusion and exclusion masks for each subject. We observed that without correction 24.4% of tracks stopped in WMH mask (Figure 2A), while with correction the percentage of tracks crossing the WMH mask was improved (13.4% *versus* 29.2%, without and with correction respectively, $p < 0.05$, Figure 2B). According to this observation, the cingulum bundle was reconstructed with a tractography pipeline that includes a WMH mask to limit the negative impact of hyperintensities on fiber tracking algorithms.

**Figure S1.** Principal fODF direction(s) under the lesion mask crossing the Corpus Callosum. Yellow represents the white matter hyperintensity lesion mask displayed on the corresponding T1-weighted image. In A, the peak directions extracted from the fODF are preserved and coherent under the WMH. In B, the lesions do not impact the reconstruction of the Corpus Callosum.

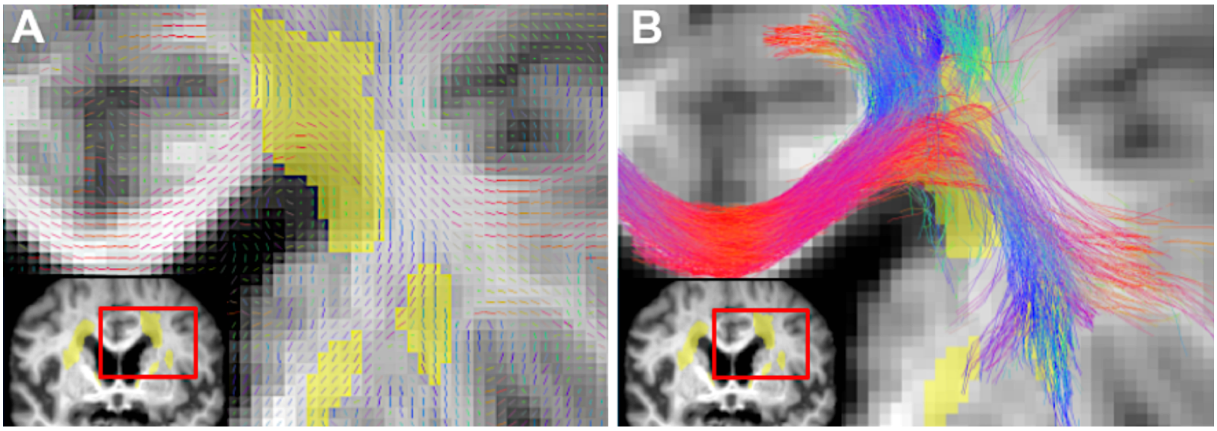


**Figure S2.** Effect of WMH mask correction on tractogram reconstruction. (A) In red tracks that stop in WMH lesion instead of grey matter and should not (B) In blue tracks that cross the WMH mask and reach grey matter regions. WMH mask is represented in yellow, both displayed on the corresponding T1-weighted image.

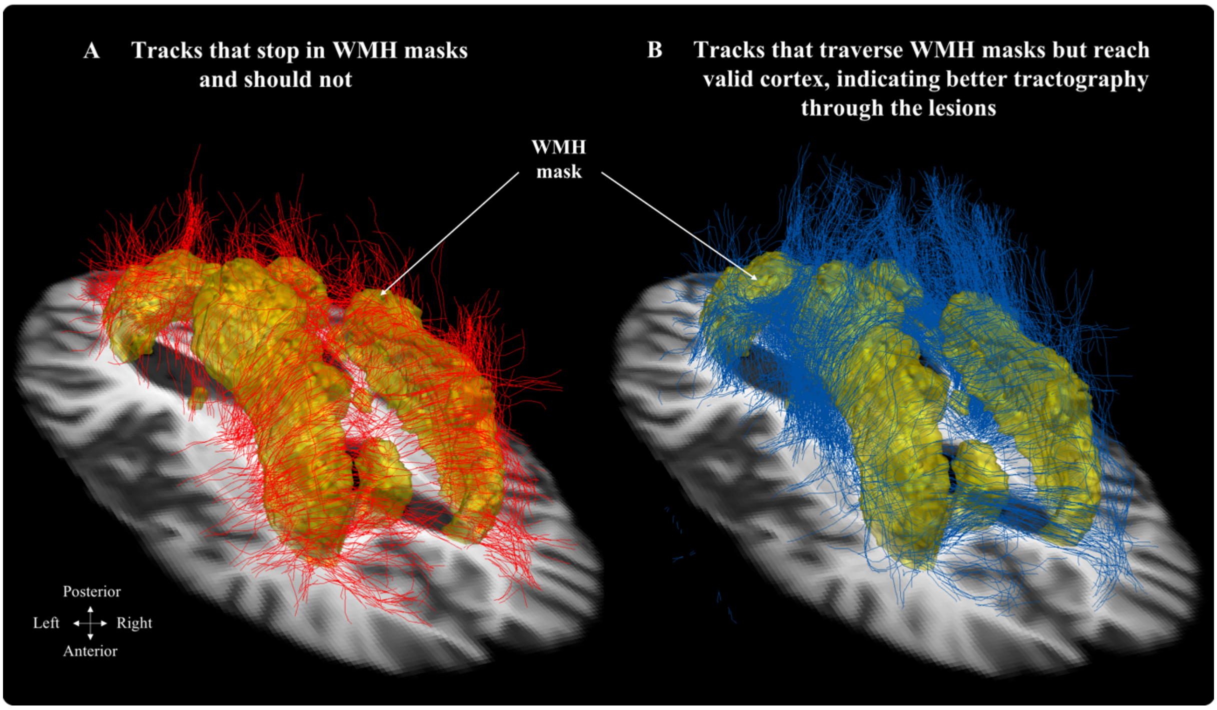

**2. Association between diffusion measures and clinical or vascular variables**

None of the cingulum diffusion measures were associated with demographic, clinical and vascular variables ($p > 0.05$, Table 1).

**Table S1.** Relationship of diffusion measures with demographic, clinical, vascular and structural variables

| | **FA** | **MD** | **AD** | **RD** | **FAt** | **MDt** | **ADt** | **RDt** | **Free water** |
|---|---|---|---|---|---|---|---|---|---|
| Age [b] | 0.107 [a] | 0.014 | 0.117 | -0.001 | 0.034 | 0.061 | 0.014 | 0.001 | -0.067 |
| | *0.504* | *0.466* | *0.209* | *0.849* | *0.668* | *0.512* | *0.551* | *0.838* | *0.501* |
| MMSE [b] | 0.175 | 0.196 | 0.166 | 0.006 | 0.191 | 0.201 | 0.178 | -0.053 | -0.183 |
| | *0.832* | *0.200* | *0.100* | *0.158* | *0.393* | *0.102* | *0.108* | *0.838* | *0.101* |
| CES-D [b] | 0.136 | 0.031 | 0.127 | -0.076 | 0.142 | 0.077 | 0.201 | -0.058 | -0.164 |
| | *0.099* | *0.458* | *0.154* | *0.402* | *0.145* | *0.379* | *0.180* | *0.226* | *0.112* |
| BMI [b] | -0.172 | -0.027 | -0.103 | 0.153 | -0.208 | -0.034 | -0.212 | -0.085 | -0.213 |
| | *0.379* | *0.827* | *0.649* | *0.486* | *0.099* | *0.828* | *0.346* | *0.250* | *0.096* |
| Gender [c] | 521 | 460 | 467 | 471 | 448 | 521 | 469 | 462 | 442 |
| | *0.807* | *0.213* | *0.301* | *0.472* | *0.348* | *0.733* | *0.297* | *0.527* | *0.172* |
| Education level [c] | 484 | 469 | 516 | 490 | 492 | 442 | 530 | 479 | 520 |
| | *0.354* | *0.789* | *0.816* | *0.582* | *0.823* | *0.401* | *0.712* | *0.504* | *0.509* |
| Diabetes [c] | 49 | 63 | 101 | 42 | 52 | 67 | 95 | 39 | 67 |
| | *0.125* | *0.442* | *0.478* | *0.113* | *0.232* | *0.369* | *0.724* | *0.157* | *0.635* |
| Hypertension [c] | 255 | 46 | 260 | 257 | 262 | 47 | 259 | 258 | 49 |
| | *0.698* | *0.176* | *0.346* | *0.352* | *0.691* | *0.224* | *0.356* | *0.562* | *0.232* |

BMI, Body Mass Index; CES-D, Center for Epidemiologic Studies-Depression scale; IST, Isaac Set Test; WMH, White Matter Hyperintensity

[a] Values are $\rho$ or U and *p-value*

[b] Spearman correlation, $\rho$ (rho)

[c] Mann–Whitney U-tests, U

**3. Association between diffusion measures and WMH volumes**

We observed a significant correlation between cingulum diffusion measures and total WMH volume ($p<0.05$; Table S2). More specifically, higher WMH volumes were associated with a higher RD (p-FDR = 0.043) using conventional DTI measures. No association was observed for cingulum WMH.

After FW-correction, higher total WMH and cingulum WMH volumes were related to higher MDt (p-FDR = 0.04 and 0.038, for total and cingulum WMH volumes respectively) and RDt (p-FDR = 0.033 and 0.029, respectively) along the cingulum bundle. No association was observed between free water content and WMH volumes in our subjects.

**Table S2.** Association between diffusion measures and WMH volumes

| | DTI | | | | FW-corrected | | | | |
|---|---|---|---|---|---|---|---|---|---|
| | **FA** | **MD** | **AD** | **RD** | **FAt** | **MDt** | **ADt** | **RDt** | **Free water** |
| | $\rho$ | $\rho$ | $\rho$ | $\rho$ | $\rho$ | $\rho$ | $\rho$ | $\rho$ | $\rho$ |
| Total WMH volume (%) | -0.175 | 0.179 | 0.001 | **0.240*** | -0.194 | **0.212*** | -0.017 | **0.257*** | 0.046 |
| WMH within Cingulum (%) | -0.147 | 0.168 | 0.042 | 0.180 | -0.213 | **0.231*** | 0.010 | **0.265*** | 0.105 |

WMH, White Matter Hyperintensity volumes were expressed as % of TIV
Spearman's correlation, $\rho$
* $p < 0.05$ FDR corrected

**4. Impact of WMH within the cingulum bundle on the association between cingulum diffusion measures and changes in verbal fluency.**

Interestingly, despite the fact that only 3% of the cingulum crossed areas with WMH lesions in our population, this small WMH burden had an impact on the association between cingulum diffusion measures and changes in verbal fluency in a model adjusted for age and white matter volume of the cingulum ($p < 0.05$ FDR-corrected, Table S3). More specifically, for conventional DTI measures, lower MD was associated with IST decline at 15 seconds. No association was observed with FA, RD and AD. After FW-correction, higher MDt and RDt were associated with IST decline at 15 and 30 seconds. No association was observed with FAt and ADt. No association was observed between free water content and WMH volumes in our subjects. High free water content was associated with changes in IST score at 15 and 30 seconds.

**Table S3.** Relationship between cingulum diffusion measures and verbal fluency score (IST) before (conventional DTI) and after FW-correction

| | | DTI | | | | | |
|---|---|---|---|---|---|---|---|
| | | IST 15s | | IST 30s | | IST 60s | |
| | | *β* | $R^2$ | *β* | $R^2$ | *β* | $R^2$ |
| Model unadjusted for cingulum WMH[b] | FA | 0.003 | 0.001 | 0.04 | 0.002 | 0.061 | 0.004 |
| | MD | **-0.219*** | **0.091** | -0.24 | 0.06 | -0.178 | 0.03 |
| | AD | -0.203 | 0.089 | -0.18 | 0.03 | -0.096 | 0.009 |
| | RD | -0.146 | 0.021 | -0.147 | 0.02 | -0.144 | 0.02 |
| Model adjusted for cingulum WMH[b] | FA | 0.015 | 0.03 | 0.042 | 0.05 | 0.063 | 0.033 |
| | MD | -0.211 | 0.1 | -0.211 | 0.09 | -0.141 | 0.09 |
| | AD | -0.221 | 0.092 | -0.150 | 0.08 | -0.058 | 0.08 |
| | RD | -0.151 | 0.06 | -0.136 | 0.07 | -0.131 | 0.08 |
| | | **FW-corrected** | | | | | |
| | | IST 15s | | IST 30s | | IST 60s | |
| | | *β* | $R^2$ | *β* | $R^2$ | *β* | $R^2$ |
| Model unadjusted for cingulum WMH[a] | FAt | -0.055 | 0.003 | 0.037 | 0.001 | 0.049 | 0.002 |
| | MDt | **-0.368*** | **0.14** | **-0.308*** | **0.1** | -0.226 | 0.06 |
| | ADt | -0.116 | 0.013 | -0.211 | 0.05 | -0.125 | 0.016 |
| | RDt | **-0.333*** | **0.12** | -0.162 | 0.03 | -0.148 | 0.022 |
| | FW | **-0.353*** | **0.13** | **-0.241*** | **0.1** | -0.126 | 0.2 |
| Model adjusted for cingulum WMH[b] | FAt | -0.064 | 0.04 | 0.013 | 0.06 | 0.023 | 0.07 |
| | MDt | **-0.365*** | **0.17** | **-0.280*** | **0.13** | -0.187 | 0.1 |
| | ADt | -0.110 | 0.05 | -0.198 | 0.09 | -0.108 | 0.09 |
| | RDt | **-0.321*** | **0.14** | -0.148 | 0.07 | -0.116 | 0.09 |
| | FW | **-0.333*** | **0.16** | **-0.226*** | **0.12** | -0.100 | 0.085 |

[a] *β*, standardized coefficient regression adjusted for age and cingulum white matter volume

[b] *β*, standardized coefficient regression adjusted for age, cingulum white matter volume and cingulum WMH volumes

$R^2$, R square value

* p < 0.05 FDR corrected
